# An integrative approach to mosquito dynamics reveals differences in people’s everyday experiences of mosquitoes

**DOI:** 10.1101/2021.09.06.459057

**Authors:** M.V. Evans, S. Bhatnagar, J.M. Drake, C.M. Murdock, S Mukherjee

## Abstract

1. Urban environments are heterogeneous landscapes of social and environmental features, with important consequences for human-nature entanglements, such as that of mosquito-borne disease.
2. Investigations into this intra-urban heterogeneity in mosquito dynamics find conflicting results, likely due to the complex socio-ecological interactions and the importance of place-based context. Integrative research, which synthesizes multiple disciplines and epistemologies, can help place ecological results into their social context to explore these place-based differences.
3. Here, we develop an integrative approach to understanding spatial patterns of mosquito burdens in urban systems by combining entomological surveys, semi-structured interviews, and sketch maps.
4. Although we found no evidence for a difference in mosquito abundance across an urban gradient, there were differences in individuals’ everyday experiences with mosquitoes. These differences were mediated by how individuals moved through public space and their vulnerability to hazards in these spaces.
5. This example of integrative research illustrates what can be gained from the inclusion of multiple epistemologies, particularly for research in socio-ecological systems.

## INTRODUCTION

Urban environments are heterogeneous landscapes of social and environmental features, with important consequences for human-nature interactions. Mosquito-borne disease transmission, for example, occurs at the intersection of specific conditions of the abiotic and biotic environment, mosquito vector, and human host. Human populations in urban areas are at particular risk for certain mosquito-borne diseases, particularly *Aedes-*borne viruses (Gubler 2011, Franklinos et al. 2019) due to high human population densities (Weaver and Reisen 2010, Rose et al. 2020), adaptation of mosquito species to urban and domestic habitats (Brown et al. 2014), the abundance of larval habitat in anthropogenic landscapes (LaDeau et al. 2015), and microclimates conducive to mosquito and virus development (Murdock et al. 2017, Wimberly et al. 2020). In fact, city-dwellers make up a substantial portion of the over 3.5 billion people at risk of vector-borne diseases globally (World Health Organization 2014). Within a city, however, mosquito-borne disease risk is uneven, a result of the underlying heterogeneity of the socio-ecological landscape (Alberti 2005, Romeo-Aznar et al. 2018)

Recently, there has been an effort to understand these spatial patterns of mosquito-borne diseases in a city, and the causes of these patterns, so that limited vector control resources can be allocated efficiently (Stone et al. 2019). The findings of these studies vary widely. Mosquito populations may be most abundant in either lower socio-economic neighborhoods (LaDeau et al. 2013, Mulligan et al. 2015) or higher socio-economic neighborhoods (Becker et al. 2014, Mulligan et al. 2015). More frequent provision of piped water can increase (Stewart-Ibarra et al. 2013, Lippi et al. 2018) or decrease (Hayden et al. 2010, Schmidt et al. 2011) mosquito populations. The effect of climate is often non-linear and dependent on regional climate trends (Misslin et al. 2016, Murdock et al. 2017, Evans et al. 2019, Wimberly et al. 2020). Clearly, many of these relationships depend on the specifics of the locale in question. Further, environmental factors (e.g. hydrology, microclimate, host distributions) are the result of and interact with the socio-political context of the city (Parham et al. 2015, Santos-Vega et al. 2016). For example, a study of water infrastructure and mosquito-borne disease in Ahmedabad, India found more malaria cases in areas with a low density of formal water infrastructure connections because informal water connections, which may not be well-maintained, led to high leakage and creation of habitat for mosquito vectors, and that these leaking connections also decreased water pressure for the whole network, influencing water storage patterns, and malaria, elsewhere in the city (Subramanian et al. 2014). Understanding mosquito-borne disease in cities may therefore benefit from a place-based, integrative approach that situates ecological findings in the social context needed to properly address the complexity of a socio-ecological system (Kinzig 2001, Mayer et al. 2006).

The study of mosquito-borne diseases is particularly plagued by a lack of interdisciplinary and integrative research, limiting the types of research questions that are typically asked and therefore the potential policy solutions considered. A review found that only 3% of the research on dengue and chikungunya, two *Aedes*-borne viruses, involves a social science approach (Reidpath et al. 2011) and the field rarely engages with critical social theory and prioritizes quantifiable metrics over qualitative analyses (King 2010, Allotey et al. 2010). In addition, critical social theory studies of mosquito-borne disease rarely include ecological or environmental pathways or data in their analysis of socio-political systems. Integrative research, ““a process by which ideas, data and information, methods, tools, concepts, and/or theories are synthesized, connected, or blended” (Repko 2012), is one methodology that combines the strengths of each of these approaches in a complementary way, without prioritizing one epistemological perspective over another (Adams 2007, MacMynowski 2007). Given the current state of the field, with its stark disciplinary boundaries, integrative research is one potential way to discover the novel research avenues needed to address mosquito-borne disease.

Our study demonstrates how an integrative methodology can build on a purely ecological approach in the context of urbanization and mosquito-borne disease in a peri-urban area in southeast Bengaluru, Karnataka, India. This region has a recent history of rapid development and is the site of the city’s outward-facing urbanization into the previously rural periphery (Verma et al. 2017). In addition, Bengaluru has experienced large outbreaks of dengue, an *Aedes-*borne virus associated with rapid urbanization, over the past decade (Balakrishnan et al. 2015). We demonstrate how the application of integrative research methods to urban ecological systems can be used to better understand people’s everyday experience with mosquitoes within the broader context of urbanization. Specifically, through this study, we address the following questions:

- How does the mosquito community’s composition and abundance shift across an urban gradient?
- What are the consequences of these changes for human-mosquito interactions?
- What additional insights can be revealed through an integrative approach that may be missed by a single discipline?

The first question is based primarily in an ecological approach, focusing on metrics of mosquito community diversity and abundance. The second question attempts to understand what these findings mean in the context of people’s interactions with mosquitoes by interpreting the results in the context of qualitative description of human-mosquito interactions. Finally, the third question reflects on what is learned by placing the results of the first two questions in the context of each other.

## METHODS

### Integrative Framework

We base our work primarily in the principles of integrative research put forth by Hirsch and Brosius (2013), but which are shared with other formulations of integrative research (e.g. Miller et al. 2008, O’Rourke et al. 2019). We embrace epistemological pluralism (e.g. multiple ways of knowing and evaluating the validity of knowledge claims) through the incorporation of quantitative and qualitative data and methodologies. Viewed through an ecological or entomological perspective, the relevant patterns are those in mosquito community diversity and abundance as they shift across an urban gradient. However, mosquito-borne disease transmission occurs at the intersection of mosquito vectors and human hosts, and so we also examine patterns in individuals’ experiences with mosquitoes, questioning where, how, and why they may encounter high mosquito burdens. The construction of nature in cities (e.g. the transformation of environmental resources into amenities) is a direct consequence of existing socio-political processes and hierarchies of power, which often reproduce social inequalities in the resulting patterns in environmental amenities across a city (Lawhon et al. 2014, Gandy 2014). By focusing on power inequalities in our analysis, we aim to identify not just the patterns in mosquito populations, but the socio-political context that shapes these patterns.

In addition to an integrative approach, we ground this study in the work of feminist scholars and geographers, who recognize the power dynamics inherent to the production of knowledge and offer alternatives to conventional scientific methods through hybrid and critical approaches to the natural sciences (Kwan 2004, Lave 2015). In practice, this means that we do not use quantitative data to validate qualitative results and do not view our own knowledge of the system as academic “experts” over that of the community members. This is especially important given the power inequality between researcher and researched and the colonial history of discounting local knowledge in the field of tropical medicine.

Our methods include traditional ecological field methods of sampling mosquito communities and calculating metrics of abundance and species diversity. However, we also attempt to place these metrics into the surrounding socio-political context through the simultaneous and interdependent consideration of qualitative data, including interviews and sketch maps. Using this mixed-methods approach, we investigate how the mosquito-human interactions change across an urban gradient as a consequence of changes to ecological and social environments. The following methods section is purposefully written in a narrative way, intertwining the entomological and social methods used, rather than presenting them as separate methods, in an effort to avoid presenting this work as a “view from nowhere” that that we are somehow neutral in the process (Haraway 1988). This is especially important given the mixed-methods approach used here because community members’ interactions with us as researchers of mosquitoes influenced our discussions with them during formal interviews. It also makes it easier to recognize our own positionality in the production of this study, particularly our positions as outsiders to the residents of Sarjapur, and recognize the knowledge produced as situated (Haraway 1988).

### Study Area and Site Selection

Sarjapur is a peri-urban town located on the southeastern periphery of Bengaluru, Karnataka, India. Although historically an agricultural area, Sarjapur is currently (as of August 2021) the site of two planned Special Economic Zones, hundred-acre campuses of two major information technology (IT) companies in the country. Currently under construction, these two developments are together projected to employ over 34,000 people (Jyothi 2012), and have spurred the development of new residential complexes in the area. Large tracts of land in the otherwise rural periphery of Sarjapur are being developed by housing developers to provide single-family homes in gated communities to the IT professionals immigrating to Sarjapur. These are planned communities whose public infrastructure (e.g. roads, pipes, vegetation) is managed by the housing developer. In the town of Sarjapur itself, high-density housing exists in the form of “sheet houses” (one-story homes with corrugated tin roofs) on the outskirts and multi-story apartment buildings in town (Fig. 1). It is in the context of this large-scale development that this study took place.

**Figure 1.**
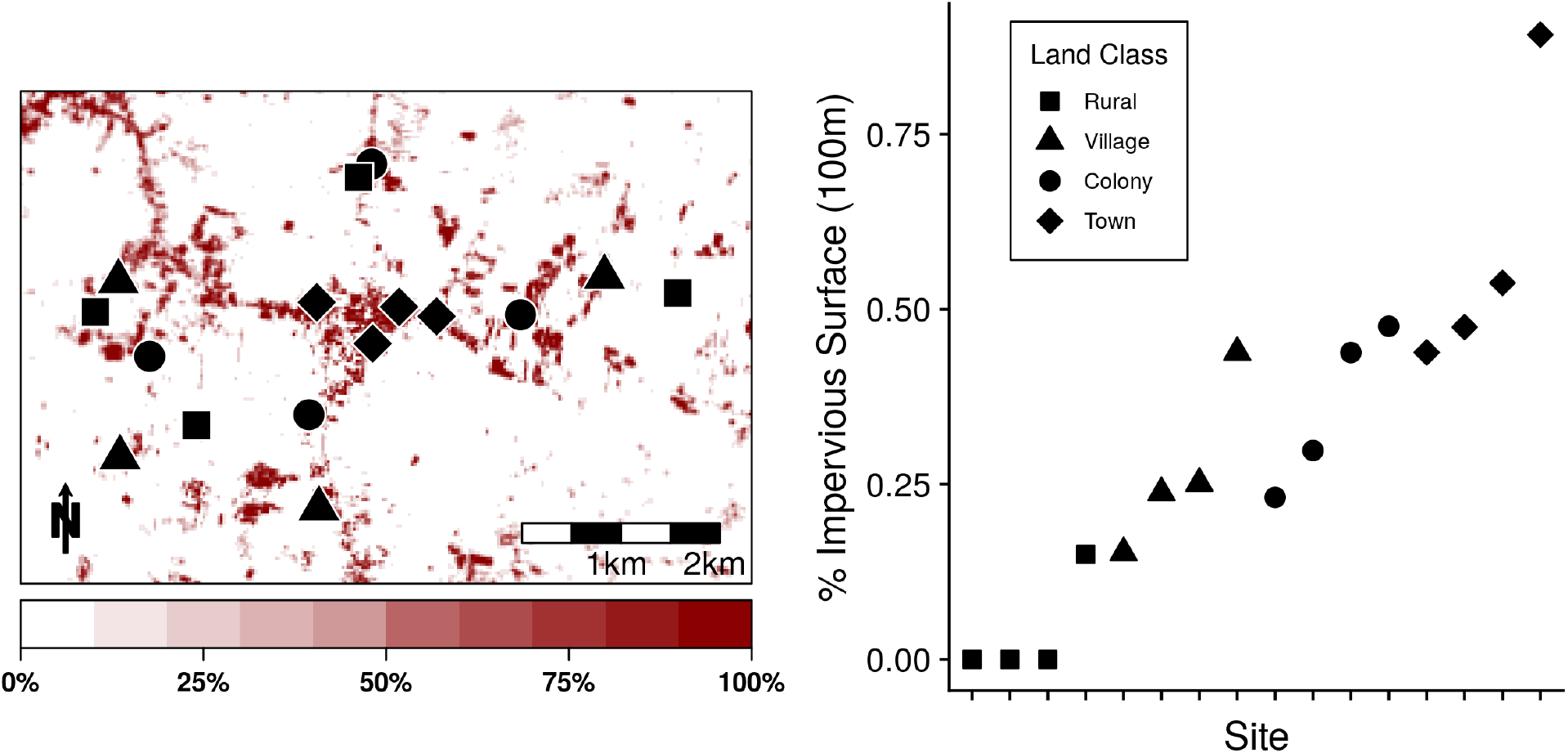
Map of sites in Sarjapur, Karnataka, India and their impervious surface values. A) Map of sites in Sarjapur, Karnataka, India. Symbols represent land classes (square: rural, triangle: village, circle: colony, and diamond: town). Color shading represents the percent of impervious surface within each 30m pixel, as illustrated on the color bar on the bottom. B) A plot of each site’s proportion of impervious surface in the surrounding 100m radius.

Within this region, we used a randomized site selection stratified across four land classes to select sixteen sites. Land classes were chosen to represent four characteristic land types of the region based on their percent of impervious surface and housing type, a qualitative measure of urbanization and an indicator of socio-economic status (Nagendra et al. 2013). Impervious surface was measured via an unsupervised land class classification of Sentinel 2 data from January - March 2019 (described in detail in Supplemental Materials S1) and approximated the amount of impervious surface in a 100m radius surrounding each site. This resulted in the following land classes: rural (low impervious surface, no housing), village (moderate impervious surface, traditional housing), town (high impervious surface, traditional housing), and colony (moderate impervious surface, Western-style housing) (Fig. 1). We divided Sarjapur into four blocks, corresponding to the East-West and North-South layout of the primary road system and associated development, and selected sites so that each block contained one site of each land class. Sites were at least 250 m from all other sites and over 1 km from sites of the same land class, except for town sites, which were clustered around the more highly developed center.

### Data Collection

Two researchers, MVE and SB, visited each site at least weekly from August to December 2019 to collect mosquito samples and conduct interviews. We visited each neighborhood regularly and took part in many formal and informal conversations throughout these four months. This allowed us to establish a presence at the site, rather than a one-time extractive sampling regime reminiscent of “parachute research”, when researchers “parachute in” to collect samples without meaningfully interacting with local community members or researchers (Lancet Global Health 2018). This familiarity with residents created a rapport that helped to recruit interview participants and conduct in-depth interviews. Weekly visits allowed us to directly observe changes to the landscape and mosquito community described in the interviews at a fine temporal scale. We conducted entomological sampling via CDC light traps and oviposition traps at the center of each site, with the permission from the resident or owner of that site. This person then served as our primary contact in the community for the interview portion of our study. As local experts, these community members also identified specific areas for trapping that were protected from disturbance and which they believed would successfully trap mosquitoes. Traps were placed in these identified areas, geolocated with a GPS device and were used as the center of each 100m site for subsequent analyses.

Oviposition traps consisted of a 1L plastic container filled with 750 mL nutrient-infused water. Nutrient infused water was prepared by mixing 20L water with 50g of ground cat food and allowing the mixture to sit for three days at room temperature. The oviposition container had two holes 1 cm from the top to allow for water to overflow. We suspended a plastic cover 10cm above the container to prevent rain and debris from entering the container. We installed two oviposition traps at approximately 1.5m height to protect traps from livestock and 5m apart at each site from August 26-28 2019, and then sampled weekly for larvae until November 11 2019, a total of 176 samples across the 16 sites. To sample, we filtered the contents of the trap through a fine-mesh filter. The water was returned to the container and we refilled the container with infused water to replace any water lost due to drying. While sampling, we often took the opportunity to discuss the mosquitoes found in that week’s collection with residents and describe the different life-stages of mosquitoes using collected samples in a clear container as an example. This provided context for our interviews with community members about mosquito habitat and its relation to water. As one community member mentioned, “I had heard of [mosquito larvae in water], but I had not seen it with my own eyes. But you both proved to us that mosquitoes can emerge from water.” All larvae and pupae were brought to the lab at Azim Premji University, where they were reared to adults, frozen at −40°C, and identified to species following (Christophers 1933, Das et al. 1990, Tyagi et al. 2014). Weekly abundances were averaged across the two oviposition traps. In instances where one trap failed during a week (e.g. tipped over or broken; 12/176 samples), we only included the working trap.

We also sampled monthly for adult mosquitoes using CDC light traps (John Hock Company and Arcturus Labs). We conducted this sampling once a month from September to November, trapping each site three times for a total of 48 trap nights. Traps were hung between 1.5 - 2 m from the ground in a covered area, to protect from rainfall. Lights were removed from the trap and traps were baited with approximately 1 kg of dry ice in an insulated plastic container hung directly next to the trap. Traps were placed in the morning and collected 24 hours later. Adult mosquitoes were frozen at - 40°C and identified to species following (Christophers 1933, Das et al. 1990, Tyagi et al. 2014). As with oviposition sampling, we often shared the results of these adult catches with residents to spark discussion of mosquito burdens. At the end of the sampling period, these results were shared with community members through the distribution of a multilingual brochure as a form of strategic communication.

Throughout the entomological sampling period, we also undertook semi-structured interviews with community members at each site, except for the rural sites which did not have houses nearby. Interviews focused on household water access and individuals’ interactions with mosquitoes. The full methods for this are described in Evans et al. (2020). Briefly, we recruited 21 community members through a combination of spatial stratification across housing types, opportunistic sampling, and snowball sampling (Stratford and Bradshaw 2016). Interviewees were adults over the age of eighteen who managed their household’s water in some capacity. Due to low literacy rates, verbal consent was obtained prior to each interview and each interview was audio recorded, translated from Kannada or Hindi, when necessary, and transcribed. Interviews consisted of a semi-structured interview and sketch mapping exercise, and generally lasted 45 minutes. The interview focused on experiences with water access and perceptions of mosquito risk in relation to water and the environment. We asked participants to identify areas with high mosquito burden on satellite map imagery, referred to as sketch maps, and discuss the environmental causes of spatial patterns of mosquitoes. In some instances, community members accompanied us to these identified areas to discuss them in more detail. This research was approved by the University of Georgia Institutional Review Board (PROJECT00000227).

### Data Analysis

We conducted analyses of the entomological data, interview transcripts, and georeferenced sketch maps simultaneously, allowing for the synthesis of observations across these different forms of knowledge. In practice, this meant pairing results from analyses of the entomological data to the transcribed interviews and sketch maps. By interpreting these results together, we noted where they did and did not agree and why. We investigated the effect of land class and impervious surface on adult mosquito diversity, adult mosquito abundance, and oviposition trap abundances using regression in a Bayesian framework that accommodated random effects and non-normal distributions (McNeish 2016). We coded interview transcripts in Atlas.ti using a narrative approach to thematic analysis, which “treats interview data as accessing various stories or narratives through which people describe their worlds” (Silverman 2003). Our coding process paid particular attention to community members’ interactions with mosquitoes and their agency and control over these interactions. Sketch maps were manually georeferenced and were analyzed alongside the interviews and entomological data using a process of grounded visualization (Knigge and Cope 2006). Results from the analysis of the entomological data and thematic analysis were interpreted in the context of finer scale spatial patterns denoted on the sketch maps and their placement in the satellite imagery depicting urbanization (e.g. maps of impervious surface and satellite imagery). We iteratively visualized sketch maps and maps of impervious surface that were linked to interview transcriptions and entomological data. If, for example, a particular area of a site was mentioned in an interview as having a characteristic that is predictive of mosquito burdens, we could connect that to the map of the site and that area’s location relative to the pattern of impervious surface at the site.

We calculated asymptotic estimates of Hill numbers from the adult mosquitoes sampled with CDC light traps for each site, aggregating across all three sampling months. Hill numbers are a set of diversity metrics that incorporate the number of species and their relative frequencies in a sample to estimate species diversity in units of “the equivalent number of equally abundant species that would be needed to give the same value of the diversity measured” (Gotelli and Chao 2013). Hill numbers are defined following Eq. 1:

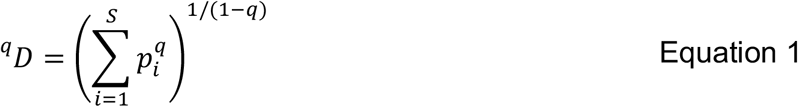

where each species *i* in the total number of species (*S)* has relative abundance *p*_*i*_ and the parameter *q*, referred to as the order, controls the sensitivity of the measure to the community’s evenness. For example, when q=0, ^*0*^*D* is equivalent to species richness, and the relative weight given to common species increases with increasing orders. For this study we calculated ^*0*^*D*, ^*1*^*D* and ^*2*^*D*, corresponding to species richness, the exponential form of Shannon entropy, and the inverse of the Simpson concentration (Chao et al. 2014a). Because the number of individuals differed greatly across sites, we used the asymptotic estimate of Hill numbers to compare these diversity metrics across these sites by extrapolating each metric to a theoretically infinite sample size following Chao et al. (2014b) using the iNEXT package (Hsieh et al. 2016) in R v. 3.6.3 (R Core Team 2018). We tested for an effect of land class and impervious surface on these metrics using generalized linear models, with the response variable transformed by subtracting 0.999 to better approximate a gamma distribution.

We used repeated-measures generalized linear mixed models to test for the effect of land class and impervious surface on adult mosquito abundance from the CDC light traps and abundance of *Aedes* mosquitoes from the oviposition traps. Because CDC light traps baited with dry ice have low-catch rates of anthropogenic *Aedes* species (Sriwichai et al. 2015), we used oviposition traps specifically to target *Aedes* abundances in addition to the CDC light traps. The abundance of the CDC light traps was the abundance of all mosquitoes caught in the trap for each site x month combination. The abundance measure from the oviposition traps represented the mean across the two traps at each site for each weekly sample, rounding up to the nearest integer.

All statistical models were implemented in Stan via the brms package (Bürkner 2017) in R v. 3.6.3 (R Core Team 2018). We tested for potential spatial autocorrelation in our response variables using the Mantel test, finding no evidence of spatial autocorrelation (Table S1). We inspected the trace plots and effective sample size and performed posterior predictive checks of each model to ensure convergence. Model formulas, priors, and sampling settings are noted in Supplemental Materials S2. Model inference was implemented by calculating the Bayes factor (BF) for a full model compared to a null model. BFs are the ratio of the likelihood of the data given the null hypothesis to the likelihood of the data given the alternative hypothesis, in this case a hypothesized relationship between a covariate (e.g. land class or impervious surface) and the response variable (Makowski et al. 2019). This metric indicates the relative evidence for one model over another, given the data, with values above 1 corresponding to more evidence for the alternative hypothesis (e.g. BF = 2 indicates the data are twice as likely to occur under the alternative hypothesis than the null hypothesis). We follow the language suggested by Jarosz and Wiley (2014) in our reporting of BFs, and purposefully choose not to provide thresholds or benchmarks for BFs in an effort to reject the arbitrary categorization of statistical measures (Wasserstein et al. 2019).

## RESULTS

### Mosquito Diversity

In the words of one community member, *“Mosquitoes, they’re everywhere. There’s nothing to avoid them*.*”* Sampling via CDC light traps caught 7,345 adult mosquitoes across 47 trap nights (one trap night of the total 48 failed), consisting of 19 mosquito species (Table 1). The majority (95.8%) of mosquitoes were *Culex quinquefasciatus*, and three sites (one village, one town, and one rural site) only had *Cx. quinquefasciatus* in their traps. Rural sites tended to have a higher species richness (^*0*^*D*) than sites in other land classes, however the 95% CI overlapped for all land classes and there was little evidence in favor of an effect of land class on mosquito species richness (^*0*^*D* BF = 0.432, Fig. 2). Although all sites had high unevenness due to the dominance of *Cx. quinquefasciatus*, there was strong evidence for an effect of land class on higher order Hill Numbers (^*1*^*D* BF = 15.71; ^*2*^*D* BF = 48.22; Fig. 2), with higher diversity at rural sites than other land classes. In comparison, there was weak evidence for a relationship between impervious surface and all order of Hill numbers at the site level (^*0*^*D* BF = 1.12; ^*1*^*D* BF = 2.27; ^*2*^*D* BF = 2.73; Fig. 2).

**Table 1.** List and count of species caught in CDC light traps over three-month sampling period.

| **Species** | **Number** | **Proportion** |
| --- | --- | --- |
| *Culex quinquefasciatus* | 7034 | 95.77 |
| *Culex gelidus* | 60 | 0.82 |
| *Aedes indicus* | 56 | 0.76 |
| *Aedes vittatus* | 50 | 0.68 |
| *Armigeres subalbatus* | 37 | 0.5 |
| *Culex vishnui* | 21 | 0.29 |
| *Culex tritaeniorhynchus* | 15 | 0.2 |
| *Aedes albopictus* | 14 | 0.19 |
| *Aedes aegypti* | 10 | 0.14 |
| *Culex psuedovishnui* | 10 | 0.14 |
| *Culex sitiens* | 10 | 0.14 |
| *Culex bitaeniorhynchus* | 9 | 0.12 |
| *Anopheles stephensi* | 7 | 0.1 |
| *Aedes spp. B* | 4 | 0.05 |
| *Anopheles subpictus* | 4 | 0.05 |
| *Aedes spp. A* | 1 | 0.01 |
| *Aedes thomsoni* | 1 | 0.01 |
| *Culex mimeticus* | 1 | 0.01 |
| *Mansonia uniformis* | 1 | 0.01 |

**Figure 2.**
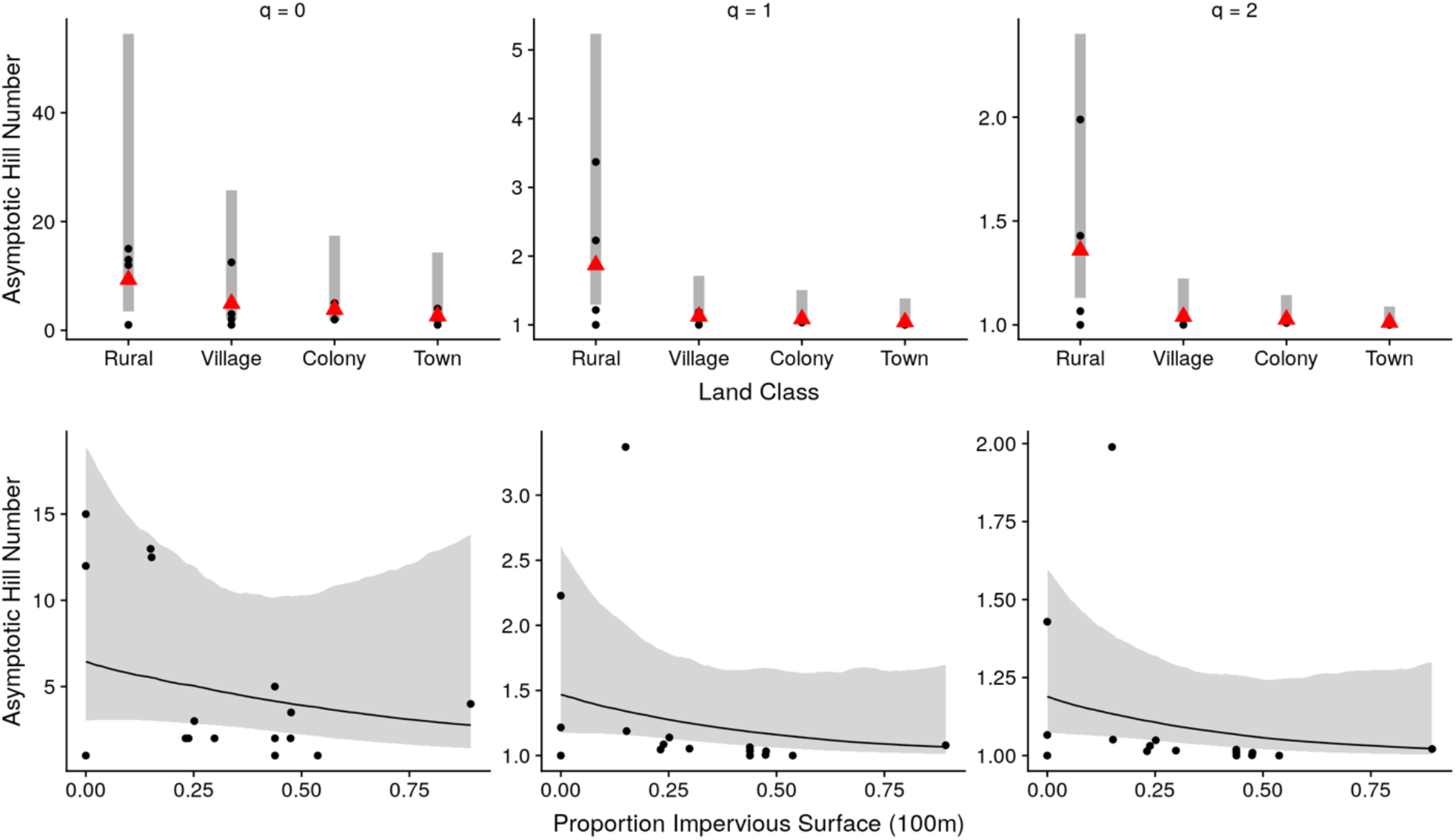
Effect of land class and impervious surface on Hill numbers. Top row: Effect of land class on asymptotic Hill numbers ^*0*^*D*, ^*1*^*D*, and ^*2*^*D*. Gray bar represents the 95% credible interval (CI) and red triangle represents the median. Raw data are plotted in black circles. Bottom row: Effect of impervious surface on asymptotic hill numbers ^*0*^*D*, ^*1*^*D*, and ^*2*^*D*. Line represents the median effect and shaded ribbon represents the 95% CI. Raw data are plotted in black circles.

Community members’ experiences of mosquito diversity tended to focus on *Aedes* mosquitoes, which are easily distinguished by their black and white markings and have been the focus of public education campaigns given their role in dengue and chikungunya transmission. However, people also distinguished between mosquito species based on their activity times, noting the difference between diurnal and crepuscular mosquitoes and how this influenced their interactions with mosquitoes. One community member described the timing of her interactions with mosquitoes, demonstrating in-depth knowledge of mosquito behaviors:

> *“The white-striped ones would be biting us around now [late afternoon]. It almost stops by the time the sun is down. The black ones bite in the night*.*”*

Some residents mentioned their habit of closing the doors and windows to their home in the early evening to keep mosquitoes from entering. *Cx. quinquefasciatus* is a crepuscular and nocturnal biter, and residents’ descriptions of their interactions with mosquitoes align with the strong dominance of *Cx. quinquefasciatus* found in the entomological surveys. For people living with mosquitoes, diversity matters because it informs the measures that people can take to avoid mosquitoes. Given that all sites were so highly dominated by the “domesticated” *Cx. quinquefasciatus*, however, there was little perception of the difference in mosquito diversity as revealed by conventional ecological metrics.

### Mosquito Abundance

We found no evidence for a difference in overall adult abundance or *Aedes* mosquito abundance, as measured by oviposition traps, across the urban gradient in Sarjapur. Statistical tests revealed no evidence for differences in adult mosquito abundance across land class (BF = 0.434, Fig. 3) or impervious surface (BF = 0.954, Fig. 3). Three *Aedes* species were present in our sites, *Aedes aegypti* (54.1%), *Aedes albopictus* (43.1%) and *Aedes vittatus* (2.73%). We also caught one *Anopheles stephensi* individual in the oviposition traps. We found little evidence for an effect of land class (BF = 0.647, Fig. S1) or impervious surface (BF = 1.202, Fig. S1) on *Aedes* abundance in oviposition traps.

**Figure 3.**
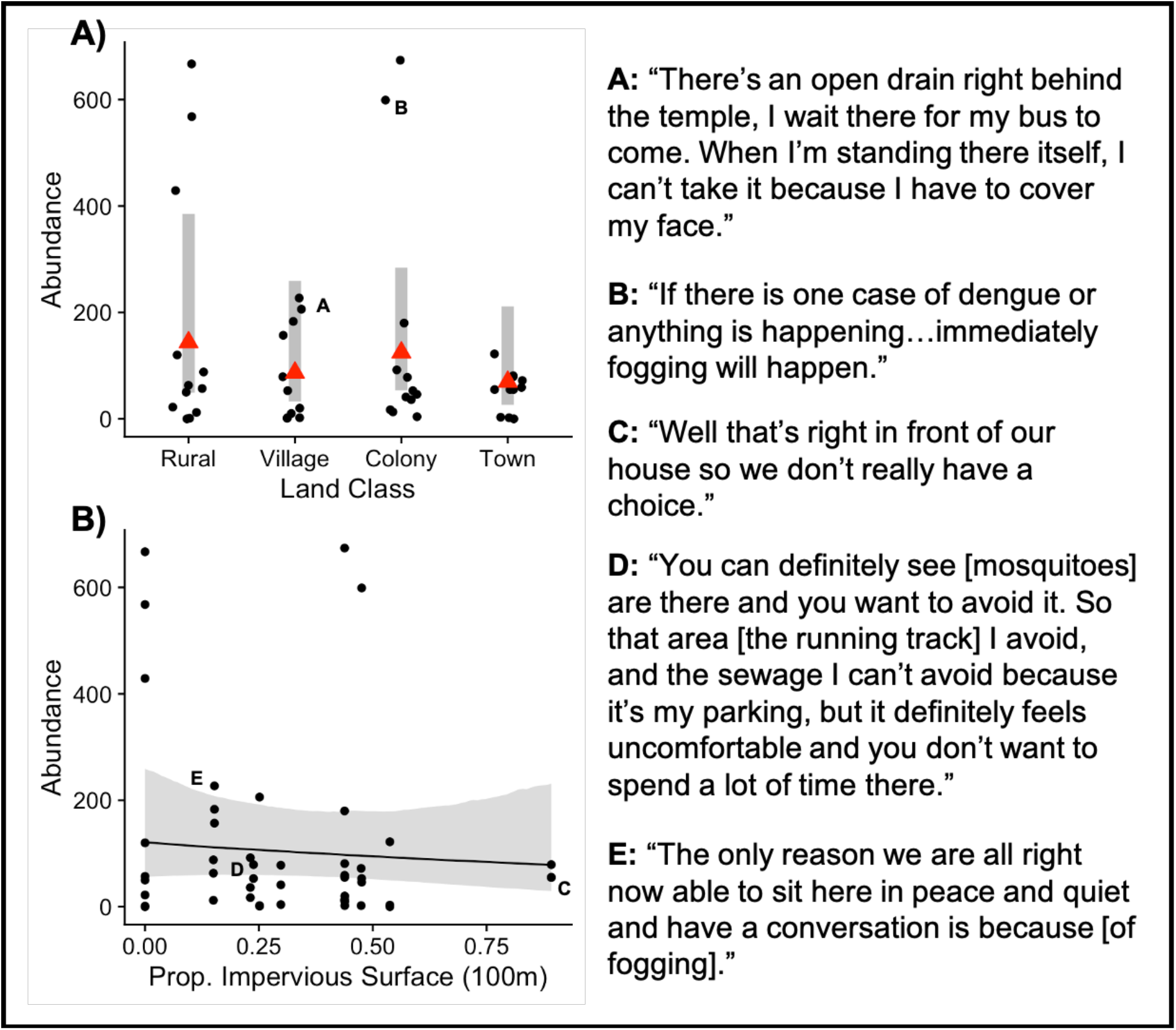
Effect of land class and impervious surface on adult abundance in CDC light traps and everyday experience with mosquitoes. A) Effect of land class on adult abundance in CDC light traps. Gray bar represents the 95% credible interval (CI) and red triangle represents the median. Raw data are plotted in black circles. B) Effect of impervious surface on adult abundance in CDC light traps. Line represents the median effect and shaded ribbon represents the 95% CI. Raw data are plotted in black circles. In both plots, points labeled with letters correspond to quotes from community members on the right-hand side of the plot.

However, peoples’ experiences with mosquitoes differed greatly across neighborhoods and did not always follow the differences in abundances that we saw from entomological sampling. One colony site had a relatively high abundance of mosquitoes compared to other sites, but the residents of this neighborhood did not express concern about the mosquito situation given their ability to request fogging by the development manager (Fig. 3). In contrast, a site in town had much lower abundances, but residents could not avoid areas with high mosquitoes, given the close proximity to their houses (Fig. 3).

In the sketch mapping exercise, community members identified areas with high mosquito burdens and described their interactions with mosquitoes in those areas. These areas included places with drainage (11/21), solid waste (13/21), or unmanaged vegetation (9/21). The size of these areas ranged from 1 - 4000 m^2^ with a median size of 72.3 m^2^. Many of these areas were close to community members’ homes and unavoidable because residents conducted everyday tasks in these outdoor spaces. One woman identified a patch of vegetation next to her home with a high burden of mosquitoes that she could not avoid:

> *“Yes, I can’t help it, I have to do my chores there. Yes, there are a lot of mosquitoes there. I have to wash the dishes there, clean my clothes also*.*”*

Further, some community members lived adjacent to open drains or waste disposal areas. One community member was dismayed by the proximity of his house to the sewage infrastructure because “the entire village’s filth comes in front of [his] house”. These narratives demonstrate that individuals differed in their ability to mitigate exposure to mosquitoes by changing their behavior or reducing mosquito abundances in public spaces through vector control.

However, these fine-scale spatial differences were obscured when we aggregated impervious surface to the site level, defined by the surrounding 100m, for use in our statistical models. When we used grounded visualization to consider the sketch maps in the context of remotely sensed impervious surface data and satellite imagery, we found that the areas identified by residents as having high mosquito burdens were areas of lower impervious surface than the surrounding landscape in 7 out of 11 sites where residents marked mosquito habitat (Fig. 4). However, there was no significant difference in mean impervious surface between the overall site mean and individually identified mosquito habitats, as determined by a paired t-test (t = 0.52522, df = 61, p-value = 0.6013). This suggests that the spatial resolution of our site-level metric (30m) was too coarse to identify these vegetative features which served as mosquito habitat within otherwise developed landscapes.

**Figure 4.**
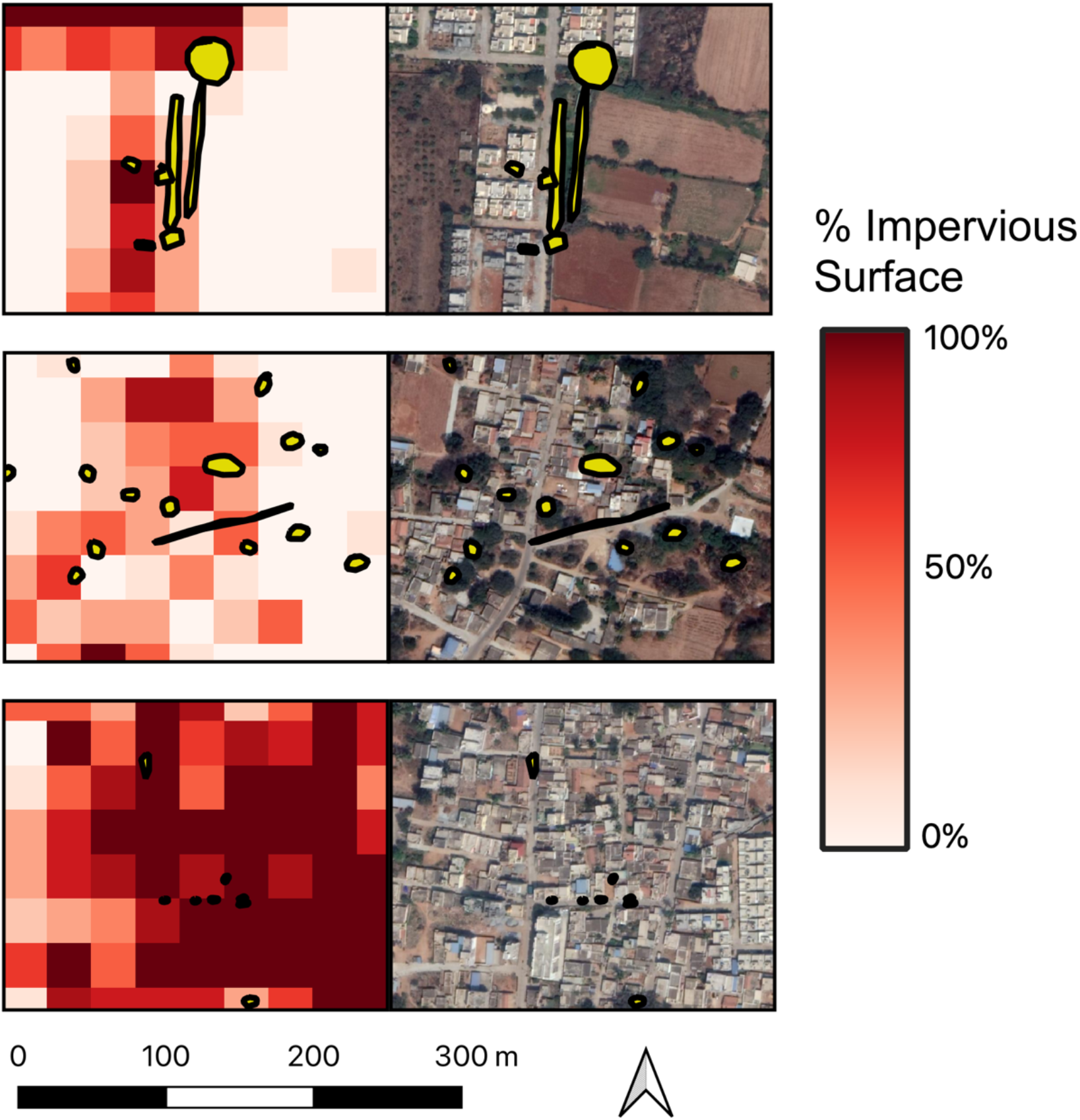
Areas of high mosquito intensity (yellow polygons) identified in sketch mapping exercise across three sites. Areas are shown over the percent impervious surface (left) and satellite imagery (right). Satellite imagery source: *Google*.

## DISCUSSION

Changes to mosquito community composition and abundances across an urban gradient is one component of intra-urban heterogeneity in mosquito-borne disease burdens. However, it is also important to understand if these changes to the mosquito community translate to changes in peoples’ experiences with mosquitoes, including both their perception of exposure and their stories of these encounters. We used an integrative approach to address these questions in parallel and a mixed-methods methodology that includes both quantitative and qualitative analyses. We found that mosquito communities were more diverse in less urbanized, rural areas, but that the dominance of all communities by *Cx. quinquefasciatus* meant that there was little perception of this difference by the population. In contrast, we found no evidence for a quantitative difference in mosquito abundance across an urban gradient, while qualitative analysis revealed that peoples’ experiences with mosquitoes was mediated by how they interacted with public spaces.

We found a difference in mosquito diversity across land classes, however this difference did not translate into differences in individuals’ experiences with mosquitoes. The mosquito community in Sarjapur was more diverse in rural land classes than in the village, town, or colony land classes, which contained a nested subset of the full community. This homogenization of the mosquito community has been seen across other urbanization or development gradients (Loaiza et al. 2017, Câmara et al. 2020) and is hypothesized to be due to changes in host composition (Goodman et al. 2018) microclimate (Townroe and Callaghan 2014), and larval habitat types (Wilke et al. 2019). However, even rural areas had high abundances of *Cx. quinquefasciatus*, which dominated the community at all land classes, and we found no evidence for a difference in total mosquito abundance across land class. Accordingly, community members perceived little difference in mosquito abundance across Sarjapur. Other studies of *Culex* species across urban gradients have found that the abundance of *Culex* species decreases with increasing urbanization (Rochlin et al. 2016, Field et al. 2019) however these effects may depend on the larger regional context (Bowden et al. 2011). Sarjapur is peri-urban, retaining characteristics of both agricultural and urbanized land classes, which may allow for similar abundances of *Culex* mosquitoes across the city.

In this study, mosquito abundance was similar across sites, but peoples’ experiences of mosquito burdens differed, primarily due to individuals’ relationships with outdoor spaces (Fig. 3). One difference was in an individual’s ability to shift their behavior to avoid interacting with mosquitoes. For upper-class households living in colonies, outdoor space was primarily used for leisure or exercise, and individuals were able to change their schedule or avoid these areas when necessary. Other households, especially those without indoor plumbing, used outdoor space for everyday domestic tasks, and their ability to shift this schedule or access another public space for this purpose was limited. These examples illustrate how individuals’ vulnerabilities differ in public spaces, drawing attention to the intersection between characteristics of the physical space and an individual’s ability to avoid or mitigate hazards of that space, such as exposure to mosquito-borne disease (Watts and Bohle 1993). Viewing disease exposure within the context of vulnerability has proven a useful approach for placing spatial patterns in infectious diseases within their social, political, and economic contexts (McLafferty 2010). Similarly, we found that individuals’ power to control a public space, via fogging or avoiding the space, helped reduce feelings of vulnerability to mosquito-borne disease exposure. Our findings regarding the everyday differences in exposure and vulnerability to mosquitoes suggest that vulnerability may be a useful lens through which to consider inequalities in mosquito-borne disease burdens as well (Chang et al. 2014).

One tension that arose through this study was the mismatch of spatial scale. Our study design focused on intra-urban heterogeneity at the scale of the city, defining each site as an area of 100m radius. However, the sketch mapping exercise revealed that people conceptualized their experience with mosquitoes on a much finer spatial scale. Between-household variation in exposure to mosquito bites can be on a magnitude of 100x difference (Guelbéogo et al. 2018), and our findings suggest that high levels of spatial variation may characterize outdoor spaces as well. Further, it has been shown that landscape heterogeneity is predictive of mosquito diversity, in addition to the dominant land class (Chaves et al. 2011). Individuals also identified patches of vegetation within a matrix of higher impervious surface as mosquito habitat, which can act as refugia for mosquitoes (Hendy et al. 2020). However, sketch mapping alone may not be able to identify all sources of mosquitoes in an area, as peoples’ movement and knowledge is constrained to areas they interact with. Top-down vector-control campaigns conceptualize patterns in mosquito burdens at coarser-levels than community members are able to, given community members’ in-depth local knowledge of the environment (Nading 2014, Dickin et al. 2014). Our discussions with community members illustrated their knowledge of mosquito dynamics in their own neighborhoods, a type of place-based knowledge that typifies a “bionomic” approach to vector control that relies on ecological relationships to identify the most productive larval habitats (Kelly and Lezaun 2013). As has been suggested elsewhere (Dongus et al. 2007, Biehler et al. 2019, Bempah et al. 2020), inclusion of local knowledge via participatory mapping and visualization activities in addition to standard entomological sampling is a promising opportunity for an alternative approach to vector control that includes place-based ecological knowledge.

Using an integrative approach allowed us to gain insight into differences that were not revealed by either method on its own. Through interviews, we expanded our conceptualization of risk from a quantity (e.g. the abundance of mosquitoes or the basic reproductive number) to one that incorporated peoples’ lived experience with mosquitoes, highlighting a difference that was otherwise missed by our entomological analysis. The collection of mosquito samples via ecological sampling created a standardized, longitudinal dataset that was easily paired with remotely-sensed imagery, to test quantitative hypotheses regarding urbanization and mosquito dynamics. The simultaneous collection of entomological and qualitative data also created opportunities for exploring the environment and everyday spaces with community members as they helped identify locations for sampling. Although these interactions occurred outside of the formal interview, we believe they enhanced our discussions with community members about the relationship between the environment and mosquitoes by providing tangible examples of these relationships, similar to other place-based interviewing methods (Holton and Riley 2014). However, an integrative approach does have additional costs associated with it, particularly in the time and expertise needed to deploy multiple methodologies. While this does not require that an integrative approach must necessarily sacrifice depth for its epistemological breadth, it does suggest that providing structural and institutional support for facilitating research across disciplines, such as through collaborative, team-based science (Leahey 2016), could reduce these costs and encourage more integrative research.

We found that mosquito community diversity was higher in rural land classes than the other three land classes, but did not find evidence for a difference in mosquito abundance across the urban gradient of Sarjapur. However, semi-structured interviews revealed a difference in how individuals experienced mosquito burdens, particularly in their vulnerability to exposure. Individuals who used outdoor space for leisure activities were able to avoid areas during peak mosquito activity periods. Those who relied on outdoor space for domestic tasks had less flexibility and often could not avoid mosquitoes, placing themselves at higher exposure to bites. Using an integrative approach shed light on the relationship between ecological indices of diversity and abundance and peoples’ everyday experience with mosquitoes, something that would not have been possible had we relied on a single discipline. A similar approach was used as the foundation of the ‘Emerging Diseases in a changing European environment’ (EDEN) project, but is rarely applied in the Global South, where urbanization rates and climate change burdens are particularly high. While we are not suggesting that all ecological studies should be integrative, we found an integrative approach to be beneficial to the study of infectious disease in urban settings.

## AUTHOR CONTRIBUTIONS

MVE, JMD, CM, and SM conceptualized the project; MVE, SB, and SM designed methodology; MVE and SB collected the data; MVE and SB analyzed the data; SM supervised the administration of the project; MVE led the writing of the manuscript. All authors contributed critically to the drafts and gave final approval for publication.

## DATA ACCESSIBILITY

Original audio files were translated and transcribed and subsequently deleted to protect the anonymity of participants. Transcriptions are saved on a secure password-protected data archive and cannot be shared without the consent of participants and a data use agreement. Geographic information, including site locations and sketch maps, are similarly protected. Entomological data and code to conduct the analyses will be deposited on figshare upon acceptance.

