## Supplemental Materials for "An integrative approach to mosquito dynamics reveals differences in people’s everyday experiences of mosquitoes"

### S1 : Unsupervised Classification of Land Classes

Impervious surface maps were created via unsupervised classification of Sentinel-2 satellite imagery in Google Earth Engine. Sentinel-2 imagery is collected globally on a 10-day frequency. We chose imagery between January - March 2019, when there is less cloud-cover, and filtered to only include individual images with less than 20% cloud cover. Because we wanted fine resolution imagery, we chose to include bands at the 10m resolution (red, blue, green, NIR). Each image was then classified into 15 land classes using the K-means Weka clustering algorithm (Frank et al. 2004). Resulting land classes were manually classified into impervious surface vs. not for each 10m x 10m pixel for each image collection date. Images were then averaged over the entire collection period, resulting in a 10m x 10m resolution image representing the uncertainty in classification. A pixel that was classified as impervious surface in at least half of the images was considered impervious surface with adequate certainty. This finalized image was then aggregated to the 30 x 30m resolution, resulting in a proportion of impervious surface in each 30m x 30m pixel.

### S2: Bayesian Model Specifications and Priors

*Hill Numbers*

The regressions for estimating the effects of land class or impervious surface on asymptotic estimates of ${}^{0}D$ used the following model formula and priors:

$$y \sim Gamma(u_i, shape)$$
$$\log(u_i) = intercept + \beta_i * x$$
$$intercept \sim Normal(0,10)$$
$$\beta \sim Normal(0,2)$$
$$shape \sim Gamma(0.01,0.01)$$

The regressions for estimating the effects of land class or impervious surface of asymptotic estimates of ${}^{1}D$ and ${}^{2}D$ used the same model formula but different priors, as the range of response variable was much smaller:

$$y \sim Gamma(\mu_i, shape)$$

$$\log(\mu_i) = intercept + \beta * x_i$$
$$intercept \sim Normal(0,2)$$
$$\beta \sim Normal(0,2)$$
$$shape \sim Gamma(0.01,0.01)$$

All models were run on three chains, each with 500 iterations following a burn-in of 500 iterations, for a total of 1500 samples.

*CDC Abundance*

The two models of CDC abundance used the following model specification and priors:

$$y \sim NegBinom(\mu_i, shape)$$
$$\log(\mu_i) = intercept_{[site]} + \beta * x_i$$
$$intercept_{[site]} \sim Normal(0,10) + \sigma_{[site]}$$
$$\sigma_{[site]} \sim Student's\ t(3,0,10)$$
$$\beta \sim Normal(0,1)$$
$$shape \sim Gamma(0.01,0.01)$$

Where *x* was either the categorical variable of land class or the proportion of impervious surface at that site and *y* was the total abundance of mosquitoes for each sampling month. All models were run on three chains, each with 1000 iterations, following a burn-in of 1000 iterations, for a total of 3000 samples.

*Oviposition Abundance*

The two models of oviposition abundance used the following model specification and priors:

$$y \sim NegBinom(\mu_i, shape)$$
$$\log(\mu_i) = intercept_{[site]} + \beta * x_i$$
$$intercept_{[site]} \sim Normal(0,10) + \sigma_{[site]}$$
$$\sigma_{[site]} \sim Student's\ t(3,0,10)$$
$$\beta \sim Normal(0,1)$$
$$shape \sim Gamma(0.01,0.01)$$

Where *x* was either the categorical variable of land class or the proportion of impervious surface at that site and *y* was the average abundance of mosquitoes per oviposition trap rounded to the nearest integer for each sampling week. All models were run on three chains, each with 1000 iterations, following a burn-in of 1000 iterations, for a total of 3000 samples.

### Supplementary Tables

| | Mantel correlation | p-value |
|---|---|---|
| Adult hill numbers | | |
| q =0 | 0.142 | 0.112 |
| q=1 | 0.136 | 0.174 |
| q=2 | 0.111 | 0.213 |
| Adult abundance | 0.065 | 0.271 |
| Oviposition abundance | 0.020 | 0.272 |

**Table S1.** Results of Mantel test for spatial autocorrelation based on 999 Monte-Carlo simulations.

### Supplementary Figures

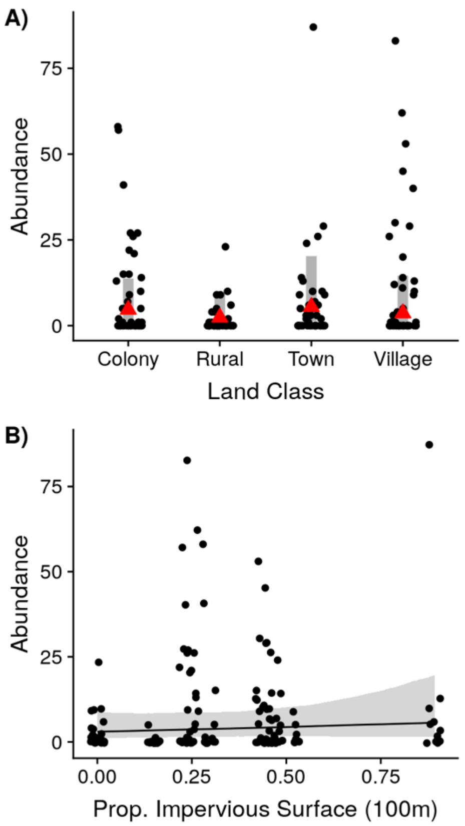


**Figure S1. Effect of land class and impervious surface on abundance from oviposition traps.** A) Effect of land class on abundance in oviposition traps. Gray bar represents the 95% credible interval (CI) and red triangle represents the median. Raw data are plotted in black circles. B) Effect of impervious surface on abundance in oviposition traps. Line represents the median effect and shaded ribbon the 95% CI. Raw data are plotted in black circles.
